# Camel urine limits proliferation and modifies cell morphology in human renal tumoral and non-tumoral cells

**DOI:** 10.1101/2022.09.12.507088

**Authors:** Carlos Iglesias Pastrana, Maria Noemi Sgobba, Francisco Javier Navas González, Juan Vicente Delgado Bermejo, Iola F. Duarte, Giovanni Lentini, Taher Kamal Sayed Osman, Lorenzo Guerra, Elena Ciani

## Abstract

The widespread ethnomedical practice of dromedary urinotherapy as a remedy against various illnesses is well recognized in traditional dromedary countries, and multiple researchers tried to unravel its bioactive potential and provide scientific evidence through *in vivo* and *in vitro* experiments. None of these studies (i) measured urine osmolarity prior to bioactivity testing, which could deeply influence the results of *in vitro* tests, nor (ii) addressed issues related to cells’ morphological changes after exposure to camel urines. Taken together, the above aspects point to the need for a “good practice” to be shared by researchers in this field, in order to reduce the variability of *in vitro* testing of camel urine bioactivity. In this work, using a set of biological samples from animals differing in sex, age, and physiological status, we investigated, the antiproliferative activity of camel urine towards human non-tumoral (HK2) and tumoral renal cells (Caki-1), through cell viability and microscopy analysis, and taking the possible influence of osmolarity into account. We employed cell lines commonly used in toxicological research which, to the best of our knowledge, have not been previously exposed to camel urine. HK2 and Caki-1 cells tolerated well mannitol-induced hyperosmolarity up to 500 mOsm/L. Significant antiproliferative effects were observed only in Caki-1 cells, when exposed to urine solutions (diluted to <500 mOsm/L) from two males out of the ten tested samples, while effects on cell morphology (elongation) were observed only in HK2 cells, when exposed to urine solutions from six samples. The significant antiproliferative effect observed only in tumoral cells looks promising for forthcoming developments in the cancer treatment field. Finally, the presented approach may serve as a guide for future research in this specific, multidisciplinary field.

## Introduction

Multiple research studies echo the popular belief within Islamic prophetic medicine that the consumption of *Camelus dromedarius* (herein after simply referred to as camel) urine, either alone or mixed with milk, is an effective and safe therapeutic remedy against various ailments [1]. Reports on the use of this animal product as a therapeutic agent in humans [2] and on its multiple bioactivities, including cytotoxic effects towards different tumor cell lines [3], inhibition of platelet aggregation [4], protection of gastric [5] and hepatic [6] epithelia, anticlastogenic activity [2] and antimicrobial properties [7], provide scientific basis in support of its putative therapeutic potential. In particular, the anticancer properties of camel urine, tested mainly *in vitro*, have harnessed increasing interest [8].

Nevertheless, some critical methodological gaps could be limiting the pharmaceutical and clinical application of the documented results. First, urine osmolarity is never measured prior to bioactivity testing and this could deeply influence the results of *in vitro* tests. Indeed, it is unknown whether the reported antiproliferative effects of camel urine could be related to the high osmolarity values (up to 3200 mOsm/L) of this animal fluid [9,10]. Additionally, the chemical composition of the urine(s) tested is hardly analyzed and this second issue hinders the identification of specific chemicals as candidates with bioactive potential [11]. Furthermore, the production of such chemicals could be influenced by multiple animal intrinsic and extrinsic factors such as sex, age, physiological status, or diet [12-14], and this variability is seldom considered. For instance, most studies have focused on the urine from female camels due to the popular belief that it is the most effective [15], thus overlooking possible gender-related effects.

In this work, we have tested the bioactivity of a set of camel urines, derived from animals of different sex, age and physiological status, while taking into account the possible influence of the samples’ osmolarity. In particular, we have assessed the antiproliferative activity of camel urine solutions towards human non-tumoral and tumoral renal cells, via microscopic examination and cell viability measurements. The two cell lines considered, HK2 and Caki-1 cells, are valuable models in toxicological research [16-22], and, to the best of our knowledge, their response to camel urine exposure is here reported for the first time.

## Materials and methods

### Urine collection and processing

All the animals included in this study are cohoused in a farm located in the Doñana National Park (southwestern Spain) and are fed on the same diet (alfalfa, beet pulp, calcium carbonate, salt, and selenium). Experimental conditions for urine sampling were adjusted following the recommendations by Emwas et al [23]. For each animal, a sterile, single use plastic bag was placed in a collection cone holder to recover mid-stream, first-morning-void urine when naturally peeing. Samples from 10 dromedary camels (Table 1) were collected on a single day in February 2020. Immediately after collection, urine was transferred to 15-mL Falcon tubes and centrifuged at 2,500 × g for 5 min at 4 °C. Once recovered, the supernatant was filtered using a 0.22 μm filter and aliquoted. For safe long-term storage, all aliquots were stored at −80° C until *in vitro* bioactivity experiments were carried out.

**Table 1.** Urine samples categorical description.

| ID | Gender | Age (years) | Physiological status |
| --- | --- | --- | --- |
| 01FD | Female | 35 | Not neutered |
| 02MF | Male | 1.5 | Bull |
| 03MH | Male | 14 | Bull |
| 04MM | Male | 28 | Neutered |
| 05MP | Male | 32 | Neutered |
| 06FP | Female | 4 | Pregnant |
| 07MR | Male | 4 | Bull |
| 08FS | Female | 19 | Not neutered |
| 09MS | Male | 30 | Neutered |
| 10MT | Male | 20 | Neutered |

### Osmolarity assessment of camel urine samples

The osmolarity of camel urine samples was measured by a VAPRO^®^ vapor pressure osmometer 5600 (Wescor, Inc., Logan, UT, USA) with fresh 290 mmol/kg, 1000 mmol/kg and 100 mmol/kg standard solutions, following the manufacturer’s instructions. A single Whatman filter paper disc (Wescor, ss-033) was placed in the central depression of the holder by metal forceps, and 10 μl of the samples diluted 1:2 with ultrapure water was expelled onto the disc. Saturated discs were rapidly transferred to the vapor pressure osmometer sample holder, and osmolarity was determined. All samples were analyzed in triplicate.

### Cell culture

The human proximal tubule epithelial cell line (HK2) and the human renal carcinoma cell line (Caki-1), proposed as a model system of proximal tubule epithelium, were kindly provided by Prof. Ciro Leonardo Pierri (University of Bari, Italy). Both cell lines were cultured in Dulbecco’s modified Eagle’s medium high glucose (Elabscience Biotechnology Inc., USA, EP-CM-L0032), supplemented with 10% foetal bovine serum (Euroclone S.p.A., Milan, Italy), in 25 cm^2^ culture flasks and were maintained at 37°C and 5% CO_2_, regularly passaged on reaching 90% confluency using a 0.25% trypsin-EDTA solution (Elabscience Biotechnology Inc., USA, EP-CM-L0446).

### Identification of osmolarity tolerance limit for cultured cells

Increasing media osmolarity conditions adjusted with mannitol were evaluated to determine the osmotic range at which no significant decrement in the viability of cells is produced. Mannitol is a cell membrane-impermeable, non-metabolizable sugar [24], used as a reference osmolyte to study tolerance limits of cultured cells to osmotic stress. Hyperosmolar solutions were prepared by the addition of the proper weighted powder of D (−) mannitol (Riedel-de Haën, cat.no. 33440) in the complete culture medium (whose osmolarity was 290 mOsm/L) to reach 400, 450, 500, 600, 700 and 800 mOsm/L respectively. After mannitol was dissolved, the solutions were sterilized using 0.22 μm filters. HK2 and Caki-1 cells were seeded in 96 multi-well plates at a density of 15,000 cells/well in the growth medium and allowed to attach overnight. Then, cells were incubated with hyperosmolar media adjusted with mannitol for 24, 48 and 72 hours at 37° C, 5% CO_2_. Cell viability was evaluated in the control condition (cells grown in the complete medium) and in the test conditions (cells grown in the hyperosmolar media) via a resazurin-based assay (Biotium, Inc., Fremont, CA, USA) performed according to the manufacturer’s instructions. The assay is based on the ability of living cells to reduce the oxidized non-fluorescent blue resazurin to a red fluorescent dye (resorufin) by a mitochondrial reductase. At the end of the resazurin incubation time (2 hours), the fluorescence reading was recorded at λ_Ex/Em_ 535/590nm in a FLUOstar^®^ Omega microplate reader (BMG LABTECH GmbH, Ortenberg, Germany). Hyperosmolarity conditions that showed fluorescence values not statistically different from the control condition were assumed as non-toxic.

### Urine samples preparation for in vitro testing

All the urines samples were first diluted 1:10 using the complete culture medium. Osmolarity of diluted urines was assessed as previously described, and urine samples (40%) exceeding the 500 mOsm/L value were further diluted adopting the minimum possible dilution ratio (max 1:15, final dilution ratio) necessary to keep osmolarity values below that threshold, thus trying to avoid excessive dilution of the potentially bioactive molecules present in urine samples. The 500 mOsm/L threshold was adopted as it was shown in our experiments (see the Results section) as a non-toxic osmolarity value.

### Cell viability assay

To evaluate viability of HK2 and Caki-1 cells exposed to camel urines solutions for 24, 48, and 72 hours, the resazurin assay was adopted as previously described.

### Optical microscopy analysis

The HK2 and Caki-1 cells plated and treated with hyperosmolar solutions or with camel urine-composed solutions were monitored via optical microscopy analysis, acquiring brightfield images through a Nikon Digital SIGHT camera and the NIS Elements software (version 3.00 SP7; Nikon, Turin, Italy), with 100x magnification (TE2000 inverted microscope by Nikon, Turin, Italy) to evaluate cell morphology evolution over time.

### Statistical analysis

The mean of three replicates was calculated per each cell line, treatment condition and exposure time. Since all the experimental data sets satisfied the normality assumption, a one-way analysis of variance (ANOVA) followed by Dunnett’s *post hoc* test were performed using SPSS Statistics for Windows, Version 25.0 [25]. Dunnett’s test is a multiple comparison procedure to determine whether means from each experimental group are statistically different with respect to the control group mean.

## Results

### Osmolarity of camel urine samples

Considering the strong water reabsorption capacity of camel kidneys, and the highly concentrated urines that this species is able to eliminate, the osmolarity of tested camel urine samples was measured before cell exposure. We found a great variability in the urine concentration, with all the samples showing values far from the physiological osmolarity of the extracellular environment (about 300 mOsm/L), as reported in Table 2.

**Table 2.** Measurements of camel urine samples osmolarity.

| ID | Urine osmolarity<br>(mOsm/L) |
| --- | --- |
| 01FD | 1315 |
| 02MF | 1696 |
| 03MH | 2391 |
| 04MM | 1376 |
| 05MP | 1595 |
| 06FP | 770 |
| 07MR | 730 |
| 08FS | 1328 |
| 09MS | 2060 |
| 10MT | 1859 |

### Mannitol induces a dose- and time-dependent cytotoxicity on human renal cells

A preliminary evaluation of the effect of increasing osmolarity values (above the physiological condition) on cell viability was conducted to unravel the tolerance limits of the tested cell lines (non-tumoral HK2 and tumoral Caki-1 human renal cell lines) *in in vitro* experiments. To simulate hyperosmolarity conditions, we used D (−) mannitol as organic cell-impermeable osmolyte [26], dissolved, at different concentrations, in the cell culture medium, thus increasing medium osmolarity from 290 mOsm/L (complete culture medium) up to 400, 450, 500, 600, 700, and 800 mOsm/L. Both cell lines were seeded in 96 multi-well plates and were exposed to the complete culture medium (control) and to the six different hyperosmolar solutions for 24, 48, and 72 hours, respectively. Optical microscope (100x) observation of treated and control cells was performed at each time point to evaluate the cellular morphology in response to the hyperosmolarity. As shown in Figure 1A (HK2) and 1B (Caki-1), neither cell line presented any remarkable morphological change up to the 500 mOsm/L condition, at all the time points considered. On the contrary, higher osmolarity conditions were associated with a clear decrease in the cellular volume (and cellular density), likely due to cellular death, as also suggested by the resazurin-based cell viability assay (Figure 2A and 2B, respectively).

**Figure 1.**
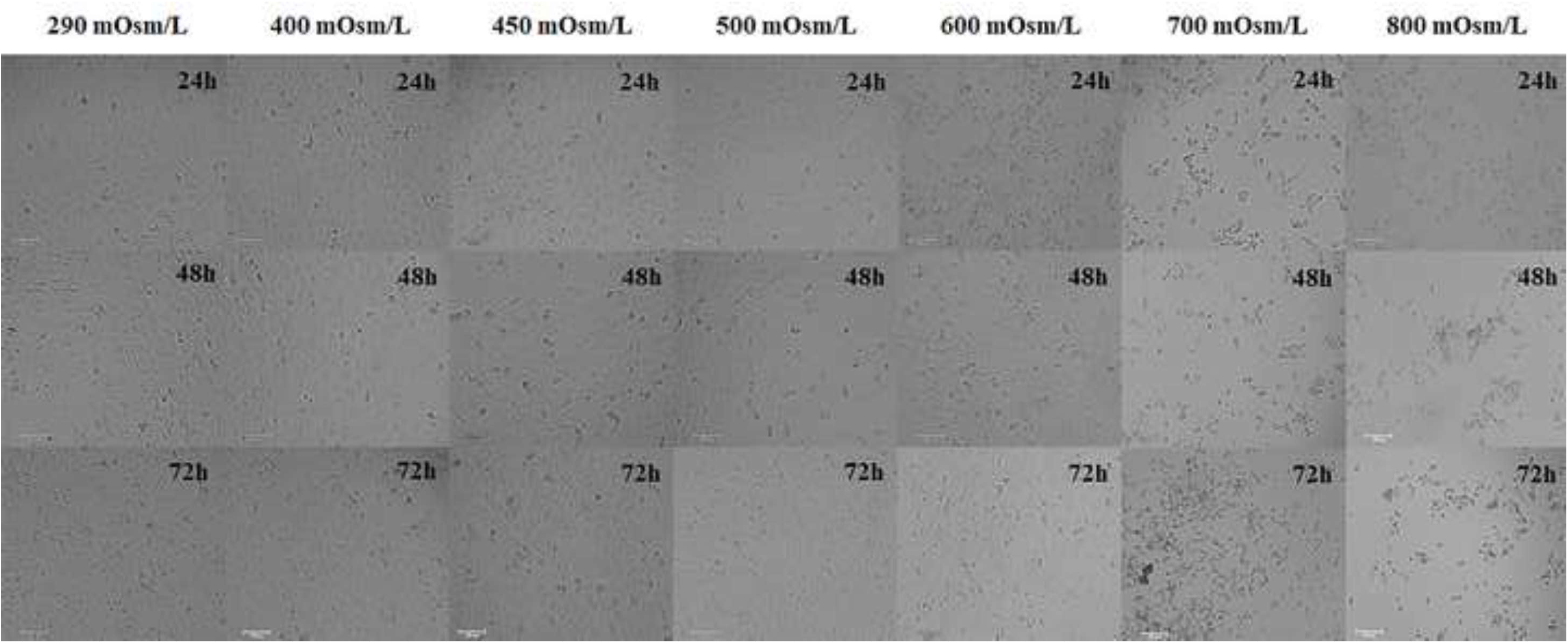
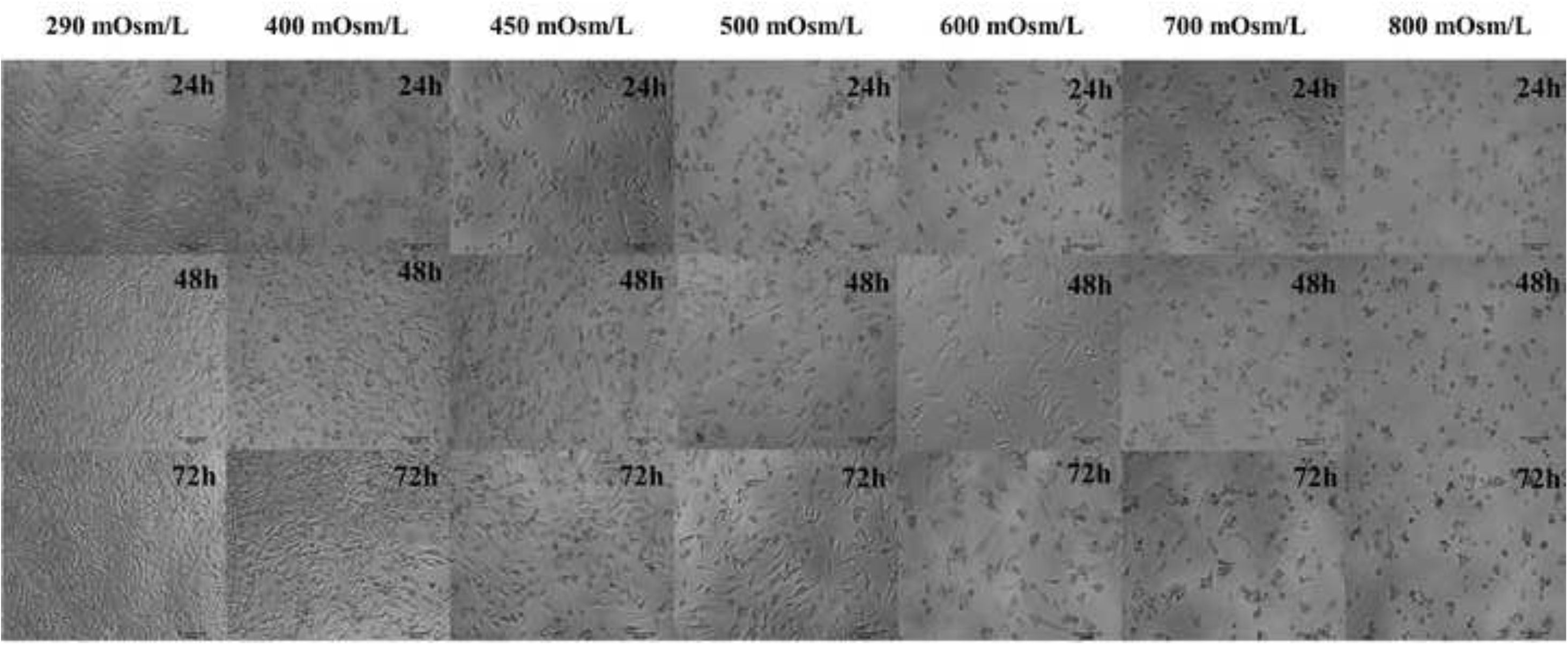
Brightfied images (100x magnification) of renal cells after exposure to hyperosmolar solutions. A) Optical microscopy analysis of human kidney (HK2) cells exposed to D (−) mannitol-composed medium. Hyperosmolarity does not significantly affect cell morphology up to 500 mOsm/L. An important and time-dependent reduction in cellular volume and cellular density was observed at 600, 700, and 800 mOsm/L. B) The human renal carcinoma (Caki-1) cell line exhibits the same behavior as HK2, when exposed to identical experimental conditions, with nearly no effect on cellular morphology and cellular density up to 500 mOsm/L. Scale bar: 100 μm.

**Figure 2.**
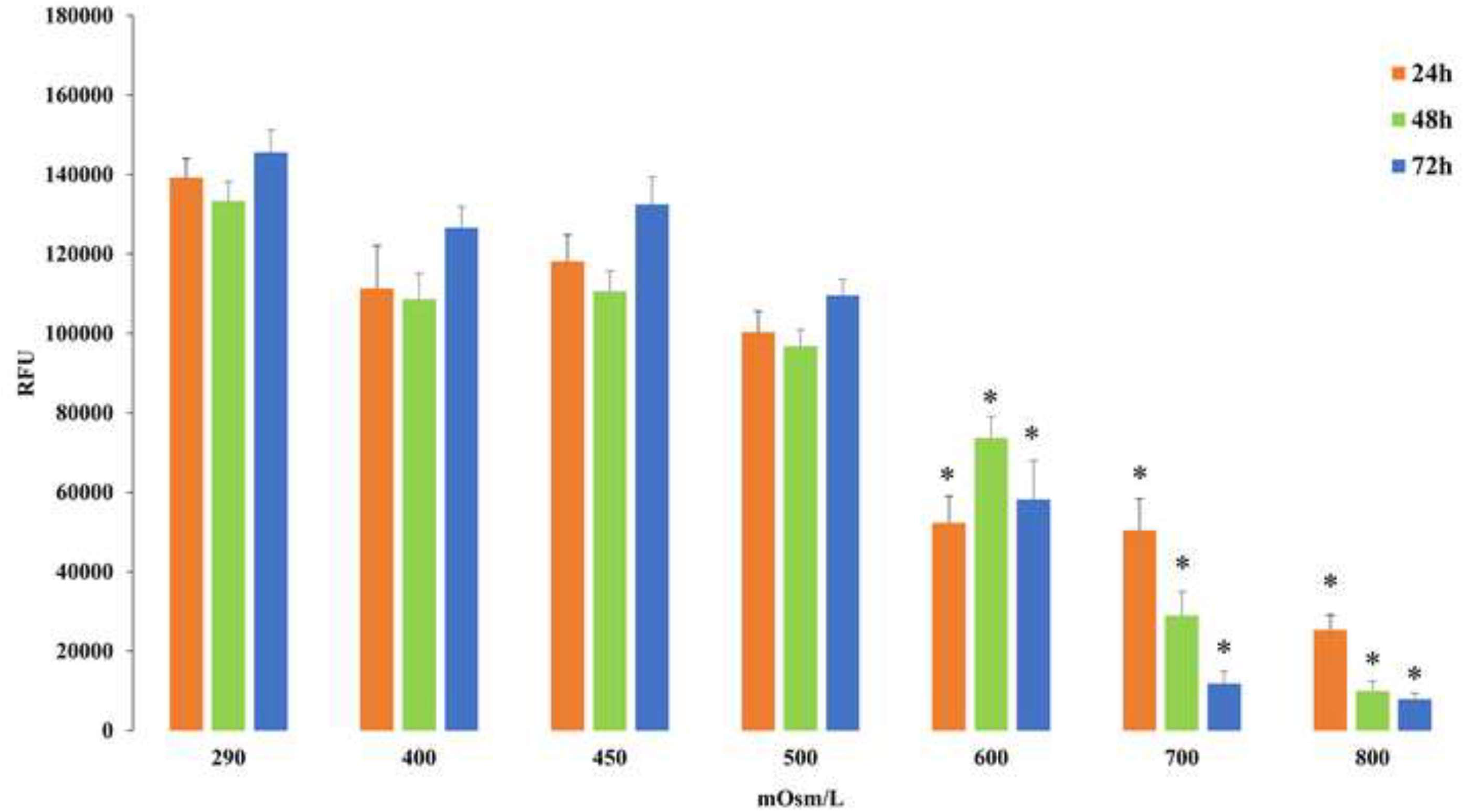
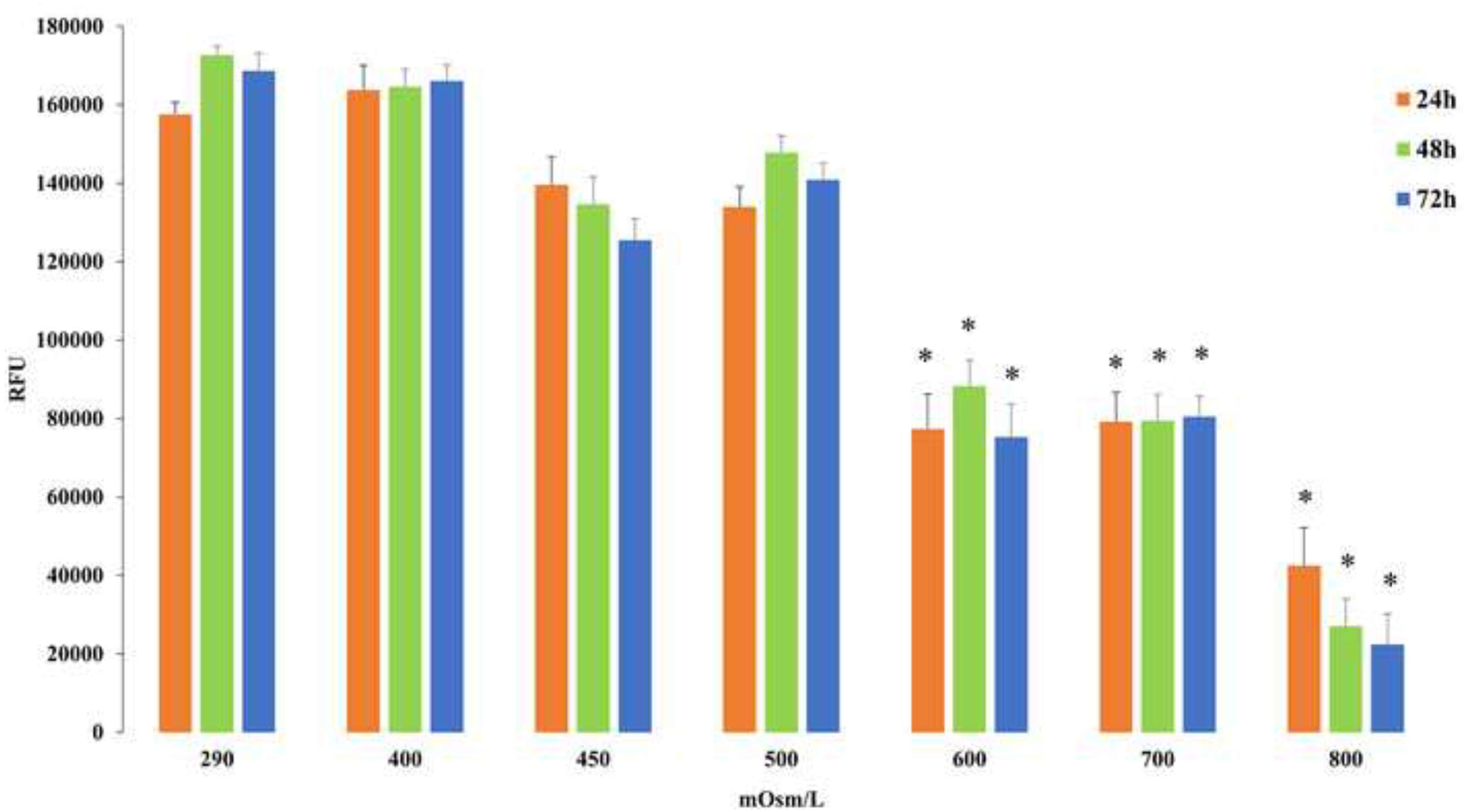
Resazurin-based cell viability assay to assess hyperosmolarity tolerance of the adopted cell lines. A) The non-tumoral HK2 cells exposed to D (−) mannitol-composed medium for 24, 48 and 72 hours exhibit a good tolerance to hyperosmolarity up to 500 mOsm/L, while 600, 700, and 800 mOsm/L cause a progressive and statistically significant decrease in cell viability (n = 9; *P < 0.05, Dunnett’s test). B) The tumoral Caki-1 cells, when exposed to identical experimental conditions as HK2, showed a slightly higher hyperosmolarity tolerance, compared to HK2 (86.7 % and 94.6 % of cell death after 72 hours of incubation, respectively, at 800 mOsm/L; *n* = 9; *P < 0.05, Dunnett’s test).

Indeed, both Caki-1 and HK2 cells showed a significant decrease in cell viability at 600, 700 and 800 mOsm/L after 24, 48 and 72 hours of incubation. In particular, a strong toxicity effect was observed at the highest concentration and the longest time of incubation, with 94.6 % and 86.7 % of HK2 and Caki-1 death cells, respectively. On the other hand, cell viability was not significantly altered, compared to control conditions, at 400, 450 and 500 mOsm/L in both our models.

### The two cell lines display dissimilar response to different urine samples

Considering the above results and the osmolarity measured for the tested urine samples (Table 2), all urines samples were diluted with cell culture medium to obtain final urine solutions with an osmolarity lower than 500 mOsm/L (see Materials and Methods). Cells were then exposed to urine solutions at the same three-time points used for the osmolarity assessments. In Caki-1 cells, for all the camel urine solutions and all the tested time points, no significant effects were detected using the resazurin-based cell viability assay, except for the urine solutions from the 02MF and 09MS camels. For these, a statistically significant viability decline was observed compared to controls at 24 hours (02MF) and 72 hours (02MF and 09MS). At the longest time of incubation (72 hours), a roughly 36% and 55% reduction in cell viability was observed for urine solutions from 02MF and 09MS compared to the control, respectively (Figure 3).

**Figure 3.**
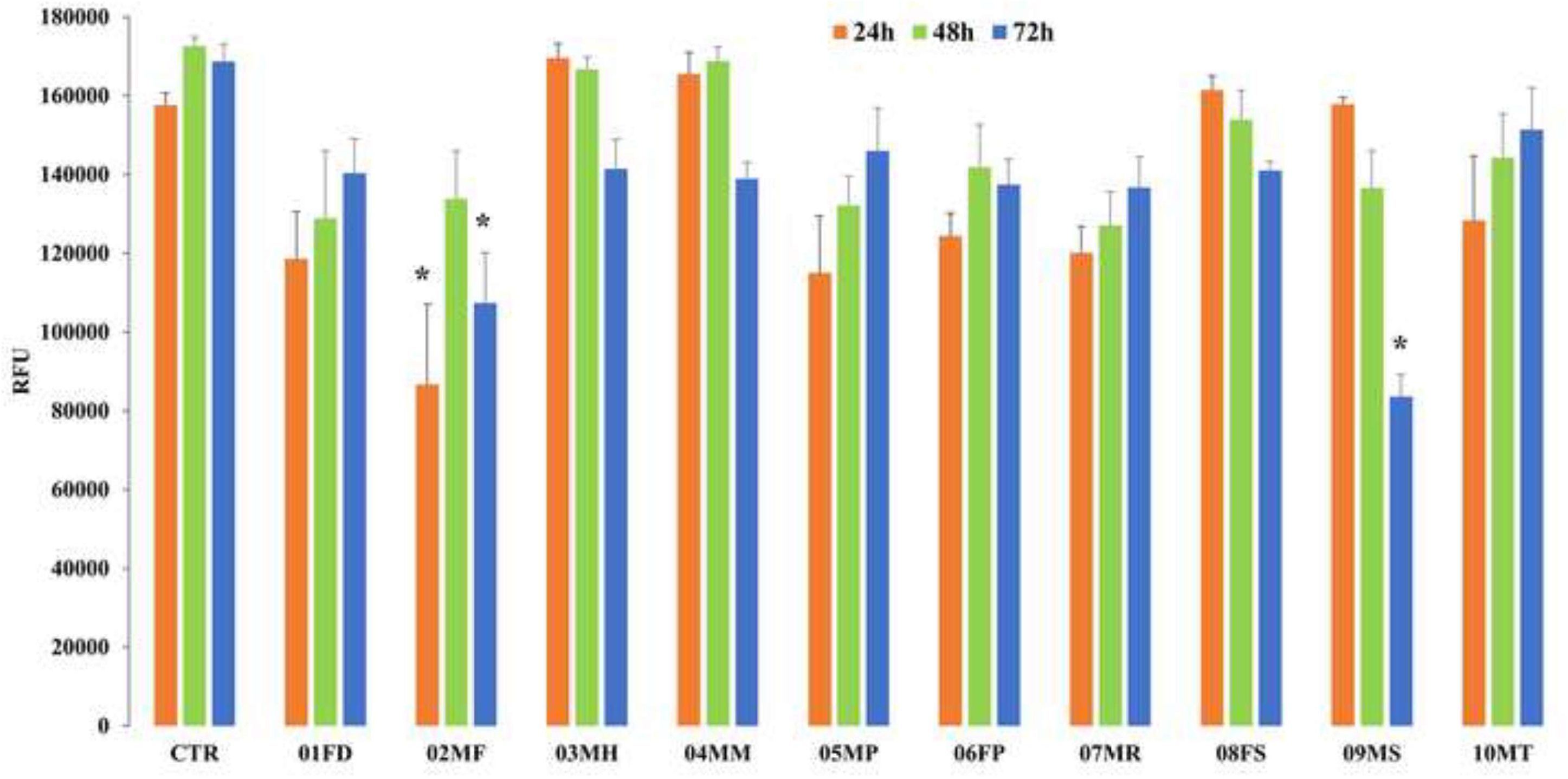
Camel urine solutions effects on tumoral renal cells viability. The tumoral Caki-1 cells were exposed to the complete culture medium (CTR) and the camel urine solutions for 24, 48, and 72 hours, and the resazurin-based cell viability assay was performed at each time point (*n* = 12). A statistically significant decrease in cell viability was observed, compared to control condition, only for solutions containing urine coming from two male camels: at 24 hours, when cells were treated with urine from 02MF (P = 0.022), and, at 72 hours, when treated with urines from 02MF (P = 0.046) and 09MS (P = 0.002). Dunnett’s *post hoc* test was applied to assess significancy of differences (*P < 0.05).

Statistically significant reduction in cell viability was not observed for the healthy cell line (HK2) in none of the urine solutions, when exposed to the same experimental conditions described above for Caki-1, although the urine solution from the 09MS camel consistently showed a marked time-dependent decreased viability (Figure 4).

**Figure 4.**
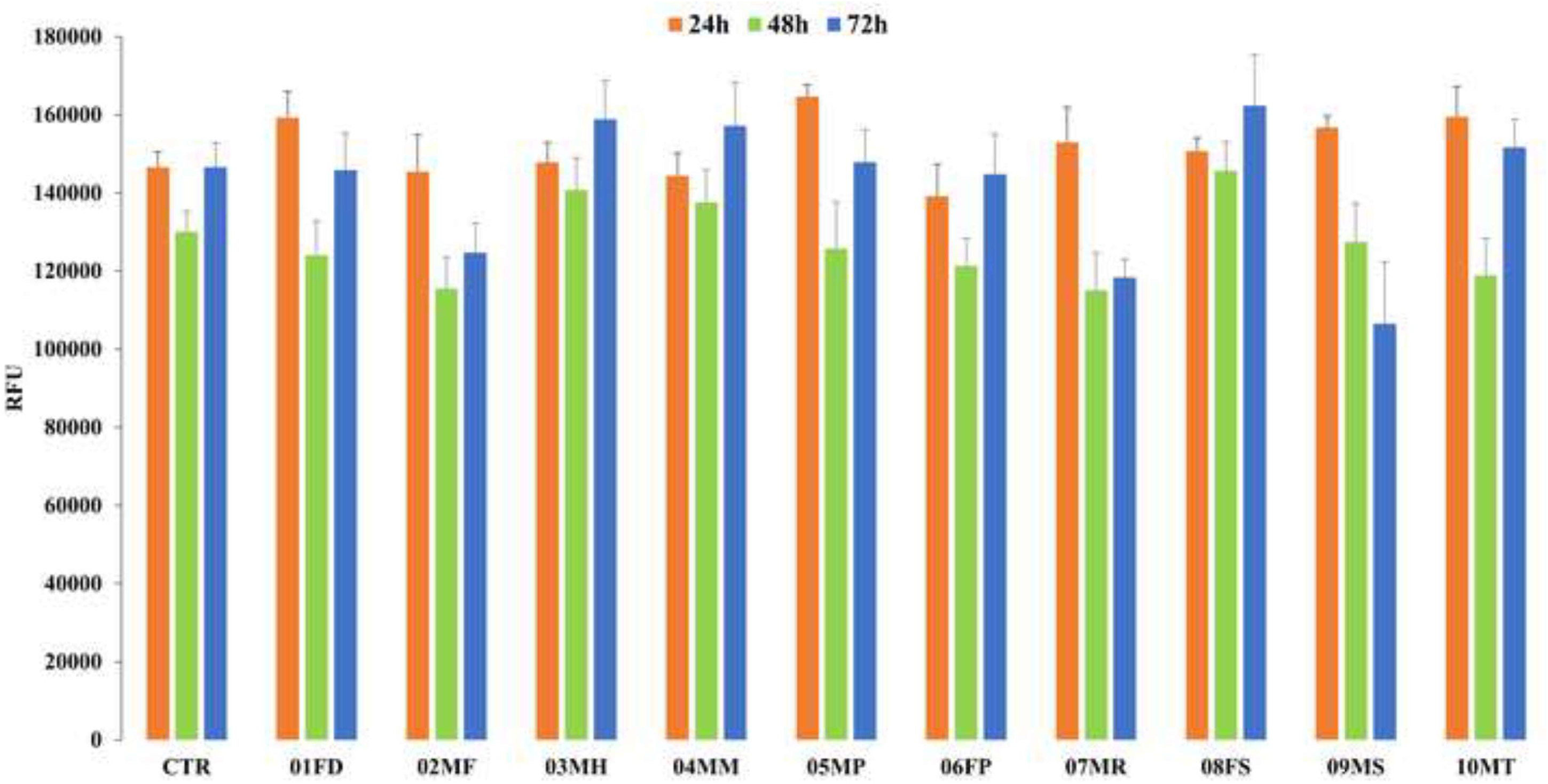
Camel urine solutions effects on non-tumoral renal cells viability. The non-tumoral HK2 cells were exposed to the complete cell culture medium (CTR) and to camel urine solutions for 24, 48, and 72 hours and the resazurin-based cell viability assay was performed at each time point (n = 12). No statistically significant decrease in cell viability was observed, compared to the control condition, applying the Dunnett’s *post hoc* test.

In what concerns cell morphology, after incubation with urine solutions, HK2 cells appeared more elongated in six out of ten samples compared to the control cobblestone shape, with the above morphology changes being gradual over time. In Figure 5, a representative image of control and treated HK2 cells, highlighting the morphological shift, is provided for the 02MF sample, while the results of the optical microscopy analysis for the remaining five samples are shown in Figures S1-S5. On the contrary, brightfield images did not show any clear modification in Caki-1 cell shape (data not shown).

**Figure 5.**
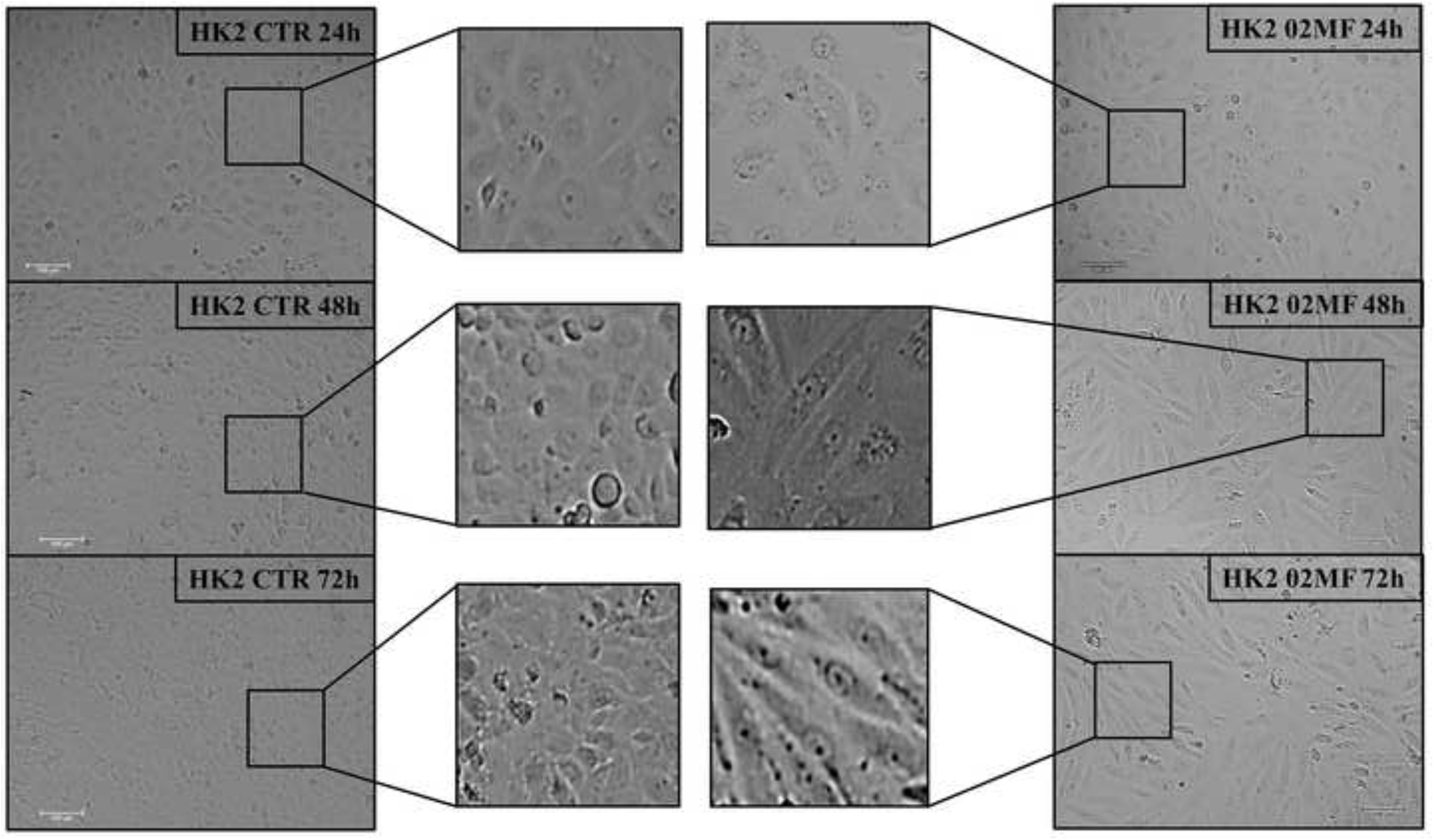
Morphological changes in non-tumoral renal cells exposed to 02MF urine solution. Morphological changes in HK2 cells, exposed for 24, 48, and 72 hours (right panel) to the solution containing urine from the 02MF sample (100x magnification). Clear and time-dependent morphological changes were observed, with HK2 becoming more elongated compared to the control (CTR) cobblestone-shaped cells (left panel). Insets of each panel represent 3x magnification of the corresponding brightfield image. Scale bar: 100 μm.

## Discussion

The currently available literature on camel urine’s bioactivity reports a plethora of potential therapeutic effects mainly referred to anticancer activity [3, 27, 28], but also to inhibition of platelet aggregation [4, 29, 30], hepatoprotective [6] and gastroprotective [5] effects, together with antibacterial [7] and antifungal [15] activities. In these studies, camel urine bioactivity has been assessed either *in vivo*, adopting mice as animal model, or *in vitro*, on human or mouse tumoral cell lines, using raw or lyophilized urines, resuspended in PBS or distilled water. In both the latter cases, the physical-chemical properties of camel urines, such as the osmolarity, that can be affected by the solvent used, have been overlooked, possibly leading to the misinterpretation of the test results. Moreover, none of these works addressed issues related with possible morphological changes of cells after exposure to camel urines, suggesting that there may be missing, though worthwhile, information. Taken together, these considerations point to the need for a “good practice” to be shared by researchers in this field, in order to reduce the variability of the *in vitro* testing of camel urine’s bioactivity.

### Osmolarity effects on human renal cells

In the camel species, the kidney presents peculiar anatomical features conferring a strong capacity of water reabsorption and the faculty to eliminate very concentrated urine, up to 3200 mOsm/L [10, 31, 32]. Urine osmolarity is a multifactorial parameter which also depends on, other than the species, the sex, and the physiological status of the animals (intrinsic factors), the geo-climatic area where camels live, their diet, access to water, and even the time of sampling during the day (extrinsic factors) [33-38]. It is well known that *in vitro* exposure to hyperosmotic stress represents a challenge for cell survival [39]. This can be particularly limiting when addressing *in vitro* studies involving effects of camel urine, as disentangling the contribution of hyperosmolarity *vs*. that ascribable to a given treatment to the observed effects on cell function could be very cumbersome. Under this scenario, we intentionally exposed our cellular models to hyperosmolarity up to 800 mOsm/L using the metabolically inert mannitol for 24, 48 and 72 hours, in order to test their tolerance to hyperosmotic stress. The HK2 (non-tumoral) and Caki-1 (tumoral) cell lines seemed to tolerate well hyperosmolarity values up to 500 mOsm/L. At higher osmolarity levels, a progressive drop in cell viability was observed, with an almost complete mortality (about 95%) at 800 mOsm/L after 72 hours of exposure. In general, the trend observed in our work is consistent with that reported by Shi et al. [40] who assessed cytotoxicity of mannitol-induced hyperosmolarity on human tubular renal cells, detecting a dose- and time-dependent effect. However, in our experimental conditions, viability levels significantly different from the control were observed at 600 mOsm/L, while in the above study viability levels significantly different from the control were already observed when 100 mM mannitol were added to their complete culture medium (roughly corresponding to 400 mOsm/L). Dissimilarity in the adopted complete culture medium composition, as well as in the number of replicates (and, consequently, in the captured biological variability, possibly impacting on statistical significance) may explain the observed differences.

### Effects of camel urine solutions on human renal cells viability and morphology

We then exposed cells to camel urines that had been diluted to keep osmolarity levels below the non-toxic threshold (500 mOsm/L), defined as per our experimental results. Interestingly, two out of the ten tested samples displayed cytotoxicity towards tumoral cells. Since all samples were adjusted to similar osmolarity, other factors, possibly related to the urines’ chemical composition, must be responsible for their bioactivity. The animals included in this study where reared under rather homogenous conditions, hence, the contribution of extrinsic factors to urine compositional variability, such as diet composition and access to water, should be minimal. In respect to intrinsic factors, our dataset included three females and seven males of varying ages and physiological status. The two bioactive samples were from two males, one being amongst the oldest neutered subjects (09MS), and the other being the youngest animal and one of three not neutered bulls (02MF). Hence, there appears to be no direct correlation between the sample bioactivity hereby examined and the animals’ age or neutering status. Another interesting aspect is that the viability of non-tumoral renal cells was not affected upon exposure to any of the tested camel urines. This is in line with other studies showing the different behavior of tumoral *vs*. non-tumoral cell lines [3, 41-44] and represents a positive result in the context of cancer treatment. On the other hand, only non-tumoral renal cells were prone to morphological changes, observed upon exposure to urine solutions from two females (01FD and 08FS) and four males (02MF, 04MM, 05MP and 09MS). Interestingly, no morphological change was detected in the mannitol-induced hyperosmolarity assays, when exposing cells to osmolarity levels below the non-toxic threshold, suggesting urine-induced cell shape modifications to be independent of osmolarity. Specific individual day-by-day management actions/events regarding individual animals, as well as constitutive (or induced) differing individual metabolic backgrounds, may contribute to the interindividual variation in the qualitative-quantitative composition of urines, which is assumed to lay behind the observed differences in cellular response. Lack of fine-scale information on the above aspects does not allow to rule out a possible contribution of cryptic variability factors. Furthermore, we cannot exclude a statistically limiting effect by the size of the sample set considered in the identification of patterns linked to the above mentioned intrinsic and extrinsic variability factors.

## Conclusions

In this study, we emphasized the need to define non-toxic osmolarity threshold levels prior to *in vitro* cytotoxicity studies when testing camel urines. The presented methodological approach may serve as a guide for future research in this specific, multidisciplinary field. We documented that, even after accounting for hyperosmolarity-induced stress, camel urine-induced effects on cellular attributes are still detectable. Noteworthy, significant antiproliferative effects were observed only in the tumoral cell line, thus suggesting room for forthcoming development in the cancer treatment field. Future studies should be devoted, on one hand, to fill the knowledge gap about camel urine composition and its variability, and, on the other hand, to deepen the studies about *in vitro* effects of single, or combined, molecules known to be present in this biological fluid, possibly prioritized through *in silico* evidence.

## Acknowledgments

The present research was carried out in the financing frameworks of the EU-funded projects CA.RA.VA.N—‘Toward a Camel Transnational Value Chain’ (Reference APCIN-2016-00011-00-00) and PRIMA (Program for Research and Innovation solutions in the Mediterranean region) CAMEL-SHIELD (Camel breeding systems: actors in the sustainable economic development of the northern Sahara territories through innovative strategies for natural resource management and marketing) project. C.I.P. is granted with a predoctoral contract (FPU Fellowship) funded by the Spanish Ministry of Science and Innovation, and M.N.S. holds a PhD fellowship granted to the University of Bari (Italy) by the Italian Ministry of University and Research (MUR), XXXVII cycle (https://www.uniba.it/it/ricerca/dottorati/37-ciclo). The authors would like to thank the personnel of the Aires Africanos farm (https://airesafricanos.com/es/) located in the Doñana National Park (southwestern Spain) for kind collaboration during urine collection. Further acknowledgment is due to the project CICECO-Aveiro Institute of Materials (UIDB/50011/2020, UIDP/50011/2020 & LA/P/0006/2020), financed by national funds through the FCT/MEC (PIDDAC).

The funders had no role in study design, data collection and analysis, decision to publish, or preparation of the manuscript.

## Author contributions

Conceptualization: L.G.

Data Curation: C.I.P.; M.N.S.

Formal Analysis: C.I.P.; M.N.S.

Funding Acquisition: E.C.

Investigation: C.I.P.; M.N.S.; L.G.

Methodology: C.I.P.; M.N.S.; L.G.

Resources: C.I.P.; L.G.; T.K.S.O

Supervision: L.G.; E.C.; F.J.N.G.; J.V.D.B.

Visualization: M.N.S.

Writing – Original Draft: C.I.P.; M.N.S.; L.G.; E.C.; G.L.

Writing – Review & Editing: C.I.P.; M.N.S.; L.G.; F.J.N.G.; J.V.D.B.; I.F.D.; G.L.; T.K.S.O.; E.C.

## Competing interests

The authors declare no conflict of interests.

## Supporting information

**Figure S1. Morphological changes in non-tumoral renal cells exposed to 01FD urine solution**. Morphological changes in HK2 cells, exposed for 24, 48, and 72 hours (right panel) to the solution containing urine from the 01FD sample (100x magnification). Clear and time-dependent morphological changes were observed, with HK2 becoming more elongated compared to the control (CTR) cobblestone-shaped cells (left panel). Insets of each panel represent 3x magnification of the corresponding brightfield image. Scale bar: 100 μm.

**Figure S2. Morphological changes in non-tumoral renal cells exposed to 04MM urine solution**. Morphological changes in HK2 cells, exposed for 24, 48, and 72 hours (right panel) to the solution containing urine from the 04MM sample (100x magnification). Clear and time-dependent morphological changes were observed, with HK2 becoming more elongated compared to the control (CTR) cobblestone-shaped cells (left panel). Insets of each panel represent 3x magnification of the corresponding brightfield image. Scale bar: 100 μm.

**Figure S3. Morphological changes in non-tumoral renal cells exposed to 05MP urine solution**. Morphological changes in HK2 cells, exposed for 24, 48, and 72 hours (right panel) to the solution containing urine from the 05MP sample (100x magnification). Clear and time-dependent morphological changes were observed, with HK2 becoming more elongated compared to the control (CTR) cobblestone-shaped cells (left panel). Insets of each panel represent 3x magnification of the corresponding brightfield image. Scale bar: 100 μm.

**Figure S4. Morphological changes in non-tumoral renal cells exposed to 08FS urine solution**. Morphological changes in HK2 cells, exposed for 24, 48, and 72 hours (right panel) to the solution containing urine from the 08FS sample (100x magnification). Clear and time-dependent morphological changes were observed, with HK2 becoming more elongated compared to the control (CTR) cobblestone-shaped cells (left panel). Insets of each panel represent 3x magnification of the corresponding brightfield image. Scale bar: 100 μm.

**Figure S5. Morphological changes in non-tumoral renal cells exposed to 09MS urine solution**. Morphological changes in HK2 cells, exposed for 24, 48, and 72 hours (right panel) to the solution containing urine from the 09MS sample (100x magnification). Clear and time-dependent morphological changes were observed, with HK2 becoming more elongated compared to the control (CTR) cobblestone-shaped cells (left panel). Insets of each panel represent 3x magnification of the corresponding brightfield image. Scale bar: 100 μm.

## Notes

### Competing Interest Statement

The authors have declared no competing interest.

