## Supplementary figures and images for "Camel urine limits proliferation and modifies cell morphology in human renal tumoral and non-tumoral cells"

### Figure S1 - 01FD.tif

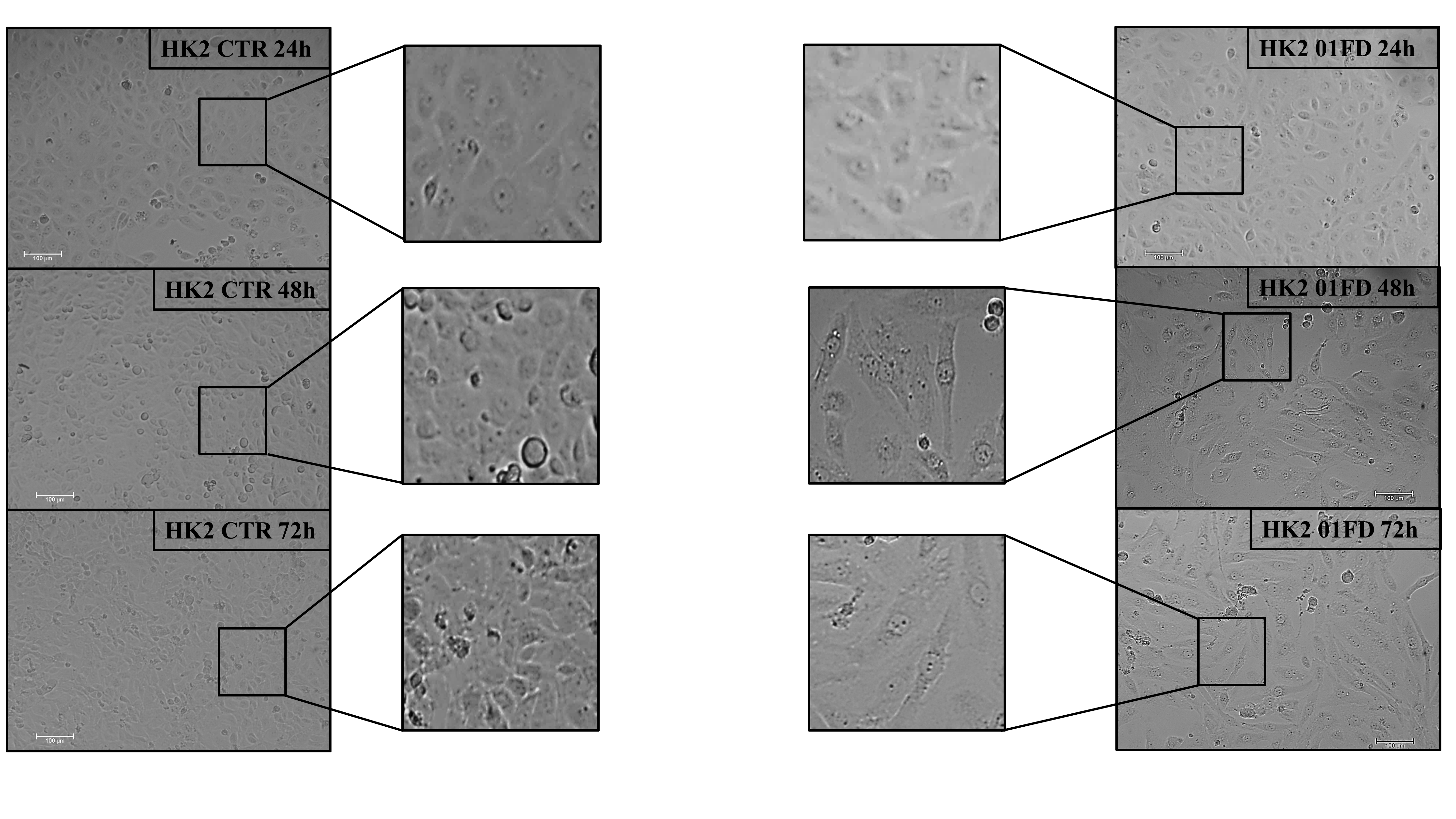

### Figure S2 - 04MM.tif

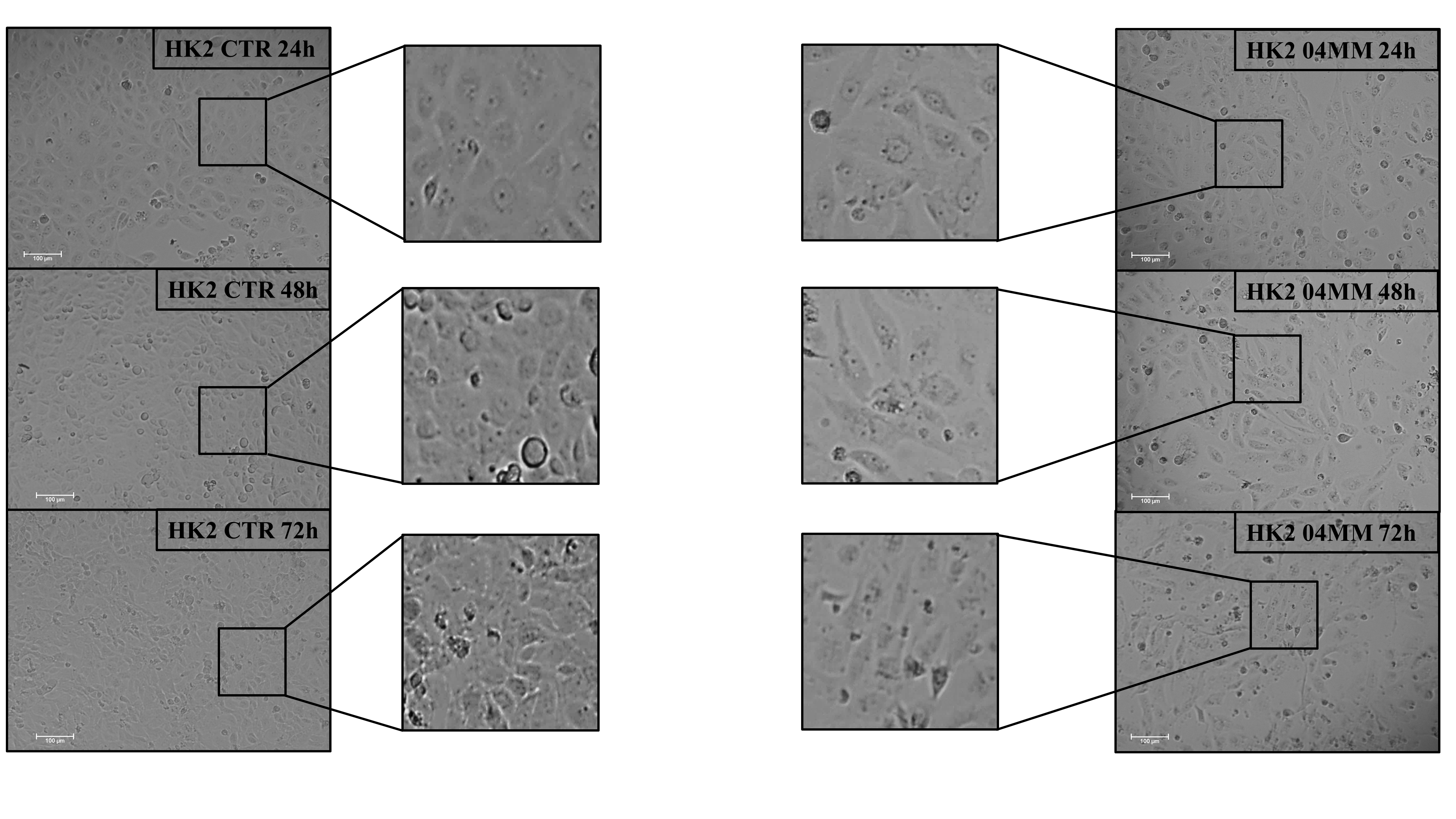

### Figure S3 - 05MP.tif

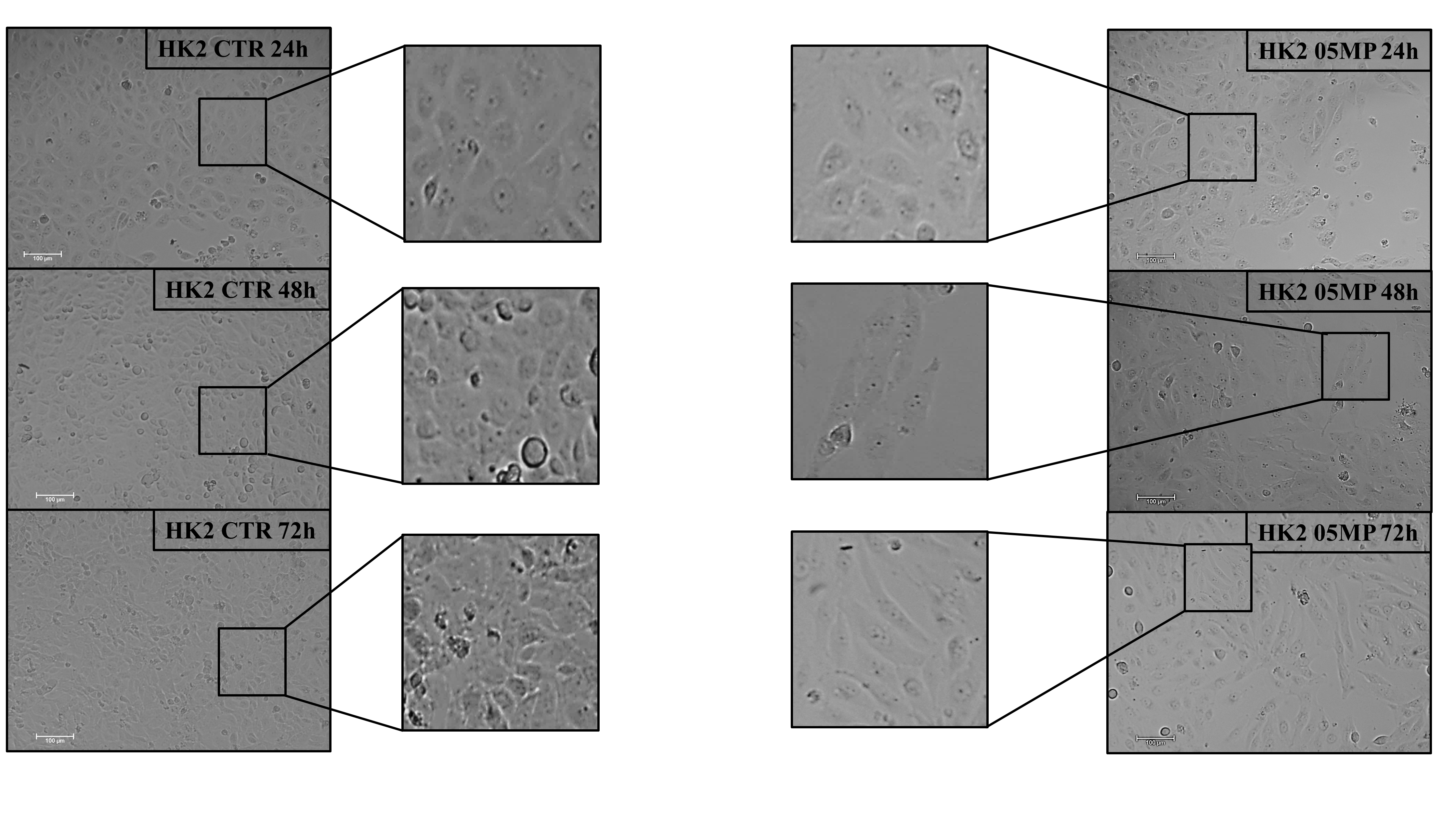

### Figure S4 - 08FS.tif

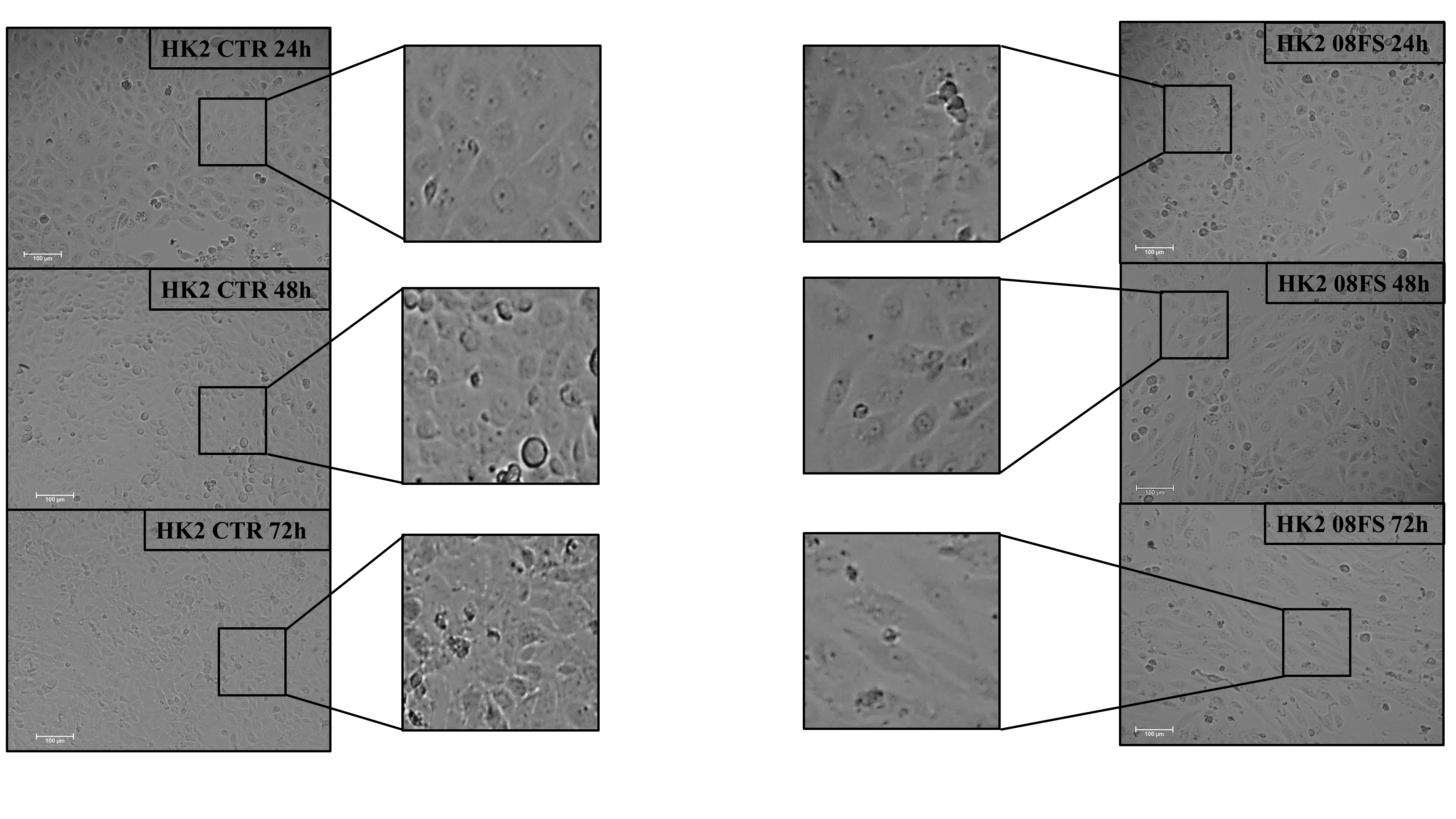

### Figure S5 - 09MS.tif

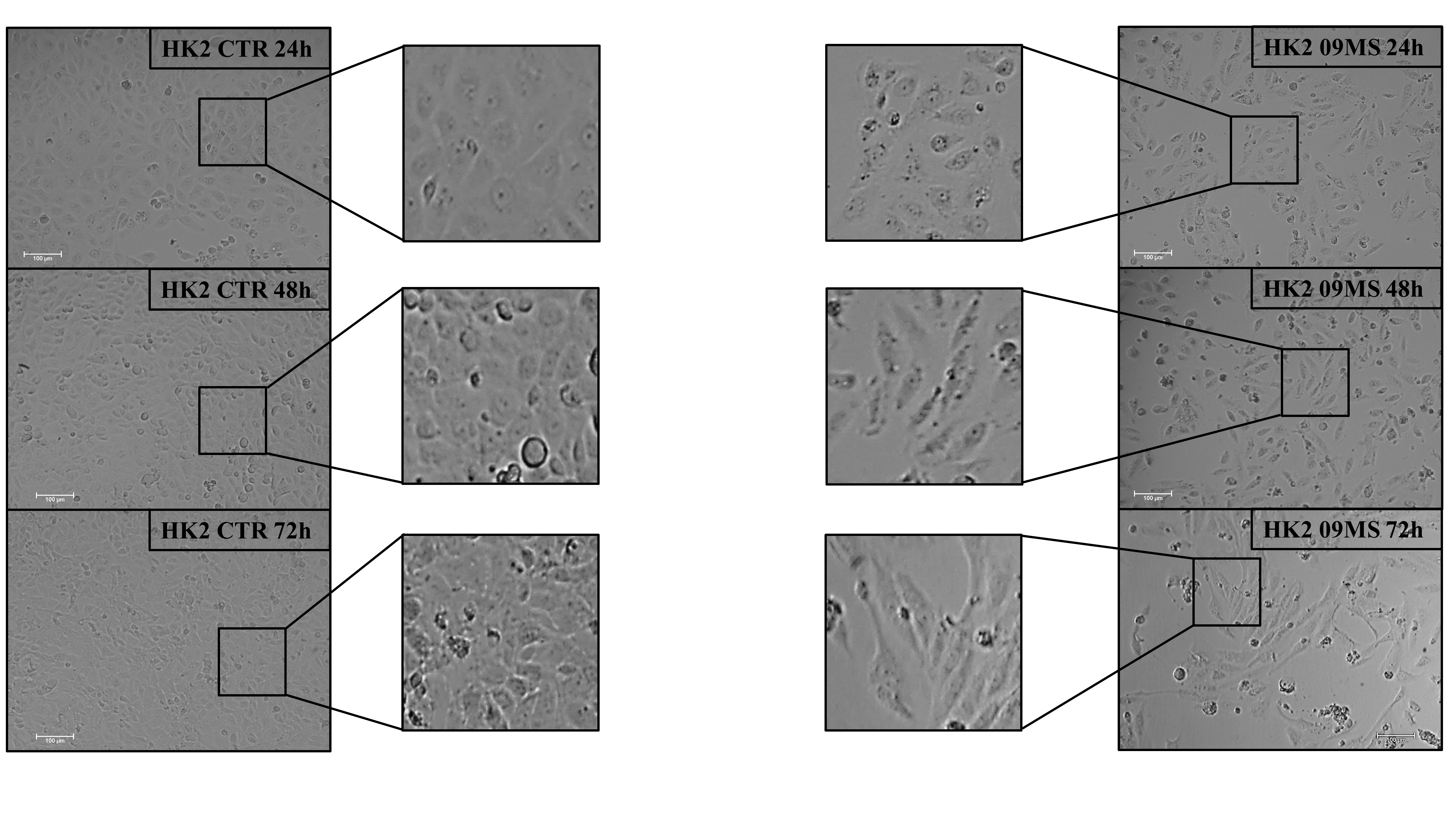
